# The organization and dynamics of corticostriatal pathways link the medial orbitofrontal cortex to future decisions

**DOI:** 10.1101/002311

**Authors:** Timothy D. Verstynen

## Abstract

Accurately making a decision in the face of incongruent options increases the efficiency of making similar congruency decisions in the future. This adaptive process is modulated by reward, suggesting that ventral corticostriatal circuits may contribute to the process of conflict adaptation. To evaluate this possibility, a group of healthy adults (N = 30) were tested using functional MRI (fMRI) while they performed a color-word Stroop task. In a conflict-related region of the medial orbitofrontal cortex (mOFC), stronger BOLD responses predicted faster response times (RTs) on the next trial. More importantly, the degree of behavioral conflict adaptation on RTs was correlated with the magnitude of mOFC-RT associations on the previous trial, but only after accounting for network-level interactions with prefrontal and striatal regions. This suggests that conflict adaptation may rely on interactions between distributed corticostriatal circuits. The convergence of white matter projections from frontal areas into the striatum was measured using diffusion weighted imaging. In these pathways, greater convergence of coticostriatal projections correlated with stronger functional mOFC-RT associations that, in turn, provided an indirect pathway that linked anatomical structure to behavior. Thus distributed corticostriatal processing may mediate the orbitofrontal cortex’s influence on behavioral updating, even in the absence of explicit rewards.

A critical feature of adaptive decision-making is the ability to modify future actions based on the success or failure of previous decisions. Consider for a moment a car spinning on a patch of ice. To stop the spin, the driver must suppress the automatic urge to turn the steering wheel against the direction of spin and instead turn the wheel into the spin. Successfully stopping the out of control car will increase the likelihood of the driver making the right decision the next time he hits a patch of ice further down the road. This rapid learning reflects a form of conflict adaptation (Gratton et al., 1992; Botvinick et al., 2001), where successful resolution of conflicting response cues (e.g., automatic desire to turn against the spin vs. correct response of turning into the spin) increases the efficiency of resolving similar conflicts in the future.

Conflict adaptation itself reflects the amalgamation of many processes, including contingency learning, feature integration, and congruency sequencing effects (Schmidt, 2013a, 2013b). Recently, reinforcement learning has been added to this list after the finding that reward signals can modulate the degree of response adaptation (van Steenbergen et al., 2009; Braem et al., 2012). Specifically, introducing financial rewards for successful resolution of conflicting cues can lead to greater adaptation on the next trial and this effect is enhanced in subjects with greater reward sensitivity (Braem et al., 2012), indicating that part of the congruency adaptation process is at least modulated by feedback signals linked to reward.

The general plurality of processes linked to conflict adaptation suggests that it relies on a broad and distributed network of brain regions. Neural studies premised on the conflict monitoring theory (Botvinick et al., 2001) usually highlight the role of the dorsal anterior cingulate cortex (dACC) and dorsolateral prefrontal cortex (DLPFC) in conflict processing. In terms of conflict adaption, neuroimaging and electrophysiological experiments have shown that signal changes in both the dACC and DLPFC are modulated on conflict trials when the previous trial also had a response conflict (Sheth et al., 2011, 2012; Kim et al., 2013a, 2013b). However, the dACC and DLPFC are not the only regions that respond to incongruent response cues. Activity in the ventrolateral prefrontal and posterior parietal cortex (Casey et al., 2000), caudate nucleus (Watanabe and Munoz, 2009), and subthalamic nucleus (Brittain et al., 2012) have all been associated with congruency processing on the current trial. It remains unclear to what extent these other regions might contribute to adaptation after a stimulus incongruency.

Taken together, the findings that reward modulates the degree of adaptation (van Steenbergen et al., 2009; Braem et al., 2012) and that basal ganglia neurons are sensitive to response conflict (Watanabe and Munoz, 2009; Brittain et al., 2012) suggest that cortico-basal ganglia systems, particularly ventral networks linked to reward processing, may be associated with trial-by-trial updating following response conflict. The current study set out to evaluate this hypothesis using a combination of structural and functional brain imaging in neurologically healthy adults while they performed a color-word version of the Stroop task (Stroop, 1935).

## Materials & Methods

### Participants

Twenty male and ten female subjects were recruited from Pittsburgh and the Army Research Laboratory in Aberdeen, Maryland. All subjects were neurologically healthy, with no history of either head trauma or neurological or psychiatric illness. Subject ages ranged from 21 to 45 years of age at the time of scanning (mean age of 31 years) and four were left-handed (2 male, 2 female). All participants signed an informed consent approved by Carnegie Mellon University and conforming with the Declaration of Helsinki and were financially remunerated for their participation.

### Stroop Task

Participants performed the color-word version of the Stroop task (Stroop, 1935; Macleod, 1991; Gratton et al., 1992; Botvinick et al., 2001) comprised of congruent, incongruent, and neutral conditions while in the MR scanner. Participants were instructed to ignore the meaning of the printed word and respond to the ink color in which the word was printed. For example, in the congruent condition, the words “RED,” “GREEN,” and “BLUE” were displayed in the ink colors red, green, and blue, respectively. In this condition, attentional demands were low because the ink color matched the prepotent response of reading the word, so response conflict was at a minimum. However, for the incongruent condition, the printed words were different from the ink color in which they were printed (e.g. the word “RED” printed in blue ink). This condition elicited conflict because responding according to the printed word would result in an incorrect response. As a result, attentional demands were high and participants needed to inhibit the prepotent response of reading the word and respond according to the ink color in which the word was printed. On the other hand, the neutral condition consisted of non-color words presented in an ink color (e.g. the word “CHAIR” printed in red ink) and had a low level of conflict and low attentional demands.

Participants were instructed to respond to the ink color in which the text appeared by pressing buttons under their index, middle and ring fingers on their right hand, each button corresponding to one of the three colors (red, green, and blue respectively) on an MR-safe response box. The task was briefly practiced in the scanner to acquaint the participant with the task and to ensure understanding of the instructions. The task began with the presentation of a fixation crosshair for 1000 ms followed by the Stroop stimulus for 2000ms during which participants were instructed to respond as quickly as possible. After the presentation of each stimulus, a fixation crosshair reappeared for an average of 4 seconds with a 40% jitter across trials to ensure accurate deconvolution of the hemodynamic response. Condition types were pseudo-randomized in an event-related fashion. A total of 120 trials were presented to each participant (42 congruent, 42 neutral, 36 incongruent). Stimuli were back-projected onto a screen located at the back of the MR bore using an MR-safe projector. Participants viewed stimuli by using a mirror attached to the top of the head coil. If necessary, vision was corrected to at least 20/40 using MR safe plastic glasses and corrective lenses.

### Behavioral Analysis

The primary behavioral variable of interest was response time (RT), recorded as the time between cue onset and registered key press (in milliseconds). All first-level analysis was restricted to correct responses. To determine condition-level effects, a one way repeated measures ANOVA was used and post-hoc one-sample t-tests were used to characterize any condition-level effects.

To measure conflict adaptation, the vector of RTs was first mean-centered. Next, all incongruent trials were selected, starting with the second trial of the series. Then these trials were categorized by the condition label of the preceding trial. Neutral trials were excluded from this comparison. Conflict adaptation was then calculated for each subject by subtracting the mean RT of Incongruent trials preceded by a congruent condition (μ_CI_) from trials preceded by an incongruent condition (μ_II_), i.e., *adaptation* = *μ_CI_* − *μ_II_*. Higher values reflect greater adaptation after repeated incongruent trials. Significance of the conflict adaptation effect was determined using a one-sample t-test on these adaptation scores.

### MRI Acquisition

All thirty participants were scanned on a Siemen’s Verio 3T system in the Scientific Imaging & Brain Research (SIBR) Center at Carnegie Mellon University using a 32-channel head coil. A high-resolution (1 mm^3^ voxel) T1-weighted brain image was acquired for all participants consisting of 176 contiguous slices using a Magnetization Prepared Rapid Gradient Echo Imaging (MPRAGE) sequence. The T1-weighted MPRAGE image was skull-stripped using a brain extraction technique (Smith, 2002) and subsequently used for registration purposes. A Blood Oxygenation Level Dependent (BOLD) contrast with Echo Planar Imaging (EPI) sequence was used for all functional MRI (TE = 20 ms; TR = 1500 ms; Flip Angle = 90°). A total of 150 T2*-weighted volumes per participant were collected for the resting state sequence, while 370 volumes were collected during the Stroop Task. Thirty contiguous slices (3.2 mm × 3.2 mm × 4 mm) were collected in an ascending and sequential fashion parallel to the anterior and posterior commissures. A 50 min, 257-direction Diffusion Spectrum Imaging (DSI) scan was collected after the fMRI sequences, using a twice-refocused spin-echo EPI sequence and multiple b-values (TR = 9,916 ms, TE = 157 ms, voxel size = 2.4 × 2.4 × 2.4 mm, FoV = 231 × 231 mm, b-max = 5,000 s/mm^2^, 51 slices). Head-movement was minimized during the image acquisition through padding supports and all subjects were confirmed to have minimal head movement during the scan prior to inclusion in the template.

### fMRI Analysis

Two participants were excluded from all analyses: one because of an error writing the data files to disk and another due to an error in the acquisition process (incorrect scan sequence used). This left a total sample size of 28 for functional imaging analysis. Functional data from each participant was processed and analyzed using the SPM8 toolbox. Prior to analysis, the EPI images for each participant were realigned to the first image in the series and corrected for differences in the slice acquisition time. All images were then coregistered to MNI-space using a non-linear spatial normalization approach (ICBM-152 space template regularization, 16 non-linear iterations). These images were then smoothed using a 4 mm isotropic Gaussian kernel.

Estimates of task-related responses at each voxel were determined using a reweighted least squares generalized linear model (GLM) approach (Diedrichsen and Shadmehr, 2005) that minimizes the influence of movement related noise in the signal. Only responses on correct trials were included in the analysis. Each trial onset was convolved with a double-gamma hemodynamic response function with each Stroop condition (Congruent, Incongruent, & Neutral) entered as a separate explanatory variable. For identification of areas with responses associated with each trial type, a condition-wise, whole-brain, random effects analysis was performed. Each Stroop condition type was estimated as a separate independent variable, providing a map of regression coefficients (β) for each condition at every voxel. To isolate incongruent trial related areas, a contrast difference between incongruent and neutral trials (β_Inc_ − β_Net_) was calculated and a one-sample t-test across all subjects was used to determine significance at each voxel. The statistical threshold for significant effects was determined using a false discovery rate (Genovese et al., 2002) across all gray matter voxels of 0.05 (q < 0.05). Clusters of more than 20 contiguously active voxels were then kept for subsequent region of interest (ROI) analyses.

After the condition-specific analysis, a single-trial event-related analysis was performed (Rissman et al., 2004). This followed the same analytical procedures as described above, with the exception that each individual trial was included as a separate independent variable in the GLM, providing a regression coefficient for each trial. The average single trial response of each ROI identified in the previous analysis was estimated by averaging the single-trial regression coefficients across all voxels in an ROI.

### Conditional Regression Analysis

Statistical mediation was performed using a conditioned regression approach and permutation-based statistical inference approach (Preacher and Hayes, 2008b) using the Bootstrap Regression Analysis of Voxelwise Observations (BRAVO) toolbox (https://sites.google.com/site/bravotoolbox). These models are designed to determine the pathways that link trial-by-trial variation in BOLD response to variation in single-trial RTs. Based on the effect sizes of single-trial BOLD estimates from previous studies (Rissman et al., 2004), as well as the estimated effect size of BOLD-RT relationships (Weissman and Carp, 2013), a minimum of 100 trials would be needed for reliable model estimation per subject (Mackinnon et al., 2002), therefore this analysis was collapsed across all three trial conditions. In addition, only ROIs with estimates of single-trial BOLD responses were correlated with both current trial RT and BOLD responses in the gyrus rectus were included as possible mediating pathways.

In the model, the vector of RTs was the dependent variable (Y), the vector of single-trial regression coefficients from the medial orbitofrontal cortex was the independent variable (X), and the single-trial regression coefficients from each of the associated ROIs were included as candidate mediating pathways (M). The selection of the independent, mediating and dependent variables were based on empirical observations (see Results). All variables were mean centered prior to analysis and pathway coefficients show in equations 1–3 were estimated using an ordinary least squares regression.

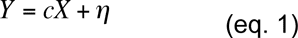

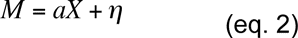

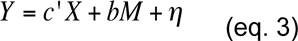

The total (*c*) pathway estimates the simple relationship between X and Y, without the inclusion of mediating variables (eq. 1). The indirect (*a*b*) pathway via each mediating variable is estimated by computing how much the X variable predicts the candidate mediator (*a* pathway, eq. 2) and the influence the mediator variable has on the Y variable (*b* pathway) when taking into account the relationship between the X and Y variables (eq. 3). Finally, the direct pathway (*c’*) reflects the residual relationship between X and Y after accounting for the influence of the mediating, indirect pathway. The η term in each equation reflects the residual noise in the estimator. This is assumed to be Gaussian and temporally independent (i.e., white noise) across trials.

A resampling-with-replacement permutation approach was used to estimate the significance of the pathways in each model. For every iteration of the algorithm, the values in the variable vectors (X, Y, and M) were scrambled independently and equations 1–3 re-estimated. The values for *a*, *b*, *c*’ and *a*b* from these permuted models were stored in a separate matrix, and this process was repeated for 1000 iterations per model. The significance of the direct and indirect paths was determined from the distribution of bootstrapped values using a bias-corrected and accelerated method (Diciccio and Efron, 1996). Statistical significance was determined after adjusting for multiple comparisons using a false-discovery rate (q) of 0.05.

### Diffusion MRI reconstruction

All DSI images were processed using q-space diffeomorphic reconstruction (Yeh and Tseng, 2011) implemented in DSI Studio (http://dsi-studio.labsolver.org). The normalization to template space was conducted using a non-linear spatial normalization approach (Ashburner and Friston, 1999) and a total of 16 iterations were used to obtain the spatial mapping function between the individual subject diffusion space, the map of QA values, to the FMRIB 1 mm template fractional anisotropy atlas. From here orientation distribution functions (ODFs) were reconstructed to spatial resolution of 2mm^3^ and a diffusion sampling length ratio of 1.25. To determine the average tractography space, a template image was generated that was composed of the average whole-brain ODF maps across all 30 subjects.

### Fiber tractography

All tractography was performed using DSI Studio (November 14^th^, 2012 build). Streamlines were generated using a generalized deterministic tractography algorithm (Yeh et al., 2013). Tractography (Fig. 5) was performed between pairs of ROI masks selected from the SRI24 Multi-Channel atlas (Rohlfing et al., 2010) and a mask of the striatum generated by merging the caudate nucleus and putamen masks and expanding this mask by 1-voxel (2 mm). Cortical targets were selected in order to encompass most frontal areas, as well as portions of the basal ganglia network. This was done by selecting 13 frontal ROI masks from the SRI24 atlas: gyrus rectus (Rectus), ventral medial prefrontal cortex (Frontal_Med_Orb), the lateral and middle orbitofrontal gyri (Frontal_Mid_Orb and Frontal_Inf_Orb), segments of the inferior frontal gyrus (operculum, Frontal_Inf_Oper; triangularus, Frontal_Inf_Tri), insula (Insula), middle frontal gyrus (Frontal_Mid), lateral superior frontal gyrus (Frontal_Sup), medial superior frontal gyrus (Frontal_Sup_Medial), precentral gyrus (Precentral), anterior cingulate (Cingulum_Ant), and the thalamus (Thalamus).

For each cortical ROI, the tracking session started with seed positions that were randomly started anywhere within the brain mask and fiber progression starting in opposite directions from a random initial orientation. Fiber progression continued with a step size of 1 mm and at each step the next directional estimate of each voxel was weighted by 20 percent of the previous moving direction and 80 percent by the incoming direction of the fiber. This continued until the underlying quantitative anisotropy (QA) index dropped below 0.15 or necessitated a turn of greater than 75-degrees. This process was repeated for 31,100,100 seeds (approximating 300 samples per voxel in the brain). Streamlines were included in the final dataset only if they met the following criteria: 1) had a length of less than 90 mm, 2) one end of the streamline terminated in the striatum mask and the other end of the streamline terminated in the cortical ROI mask.

### Structural Overlap Analysis

The main focus of the tractography analysis was to estimate convergence of cortical projections into the striatum (see Results for motivation). First the topology of projections from each cortical system to the region of task-related activity identified in the caudate nucleus was determined. For every subject and cortical ROI (see *Fiber Tractography* section), the percentage of streamlines terminating in the cluster of striatal voxels was calculated. Significance at each cortical ROI was determined using a one-sample t-test calculated across subjects, with an uncorrected threshold of p < 0.005. For simplicity of visual presentation (see Figure 5), the cortical ROIs were organized into four cluster sets: orbitofrontal (Frontal_Mid_Orb, Frontal_Inf_Orb, Rectus), medial frontal (Frontal_Med_Orb, Cingulum_Ant, Frontal_Sup_Medial), lateral frontal (Frontal_Mid, Frontal_Sup, Frontal_Inf_Oper, Frontal_Inf_Tri) and insular cortex (Insula).

To measure the amount of overlapping projections into each voxel an overlap index (OI) was calculated for each subject (*s*). This determines the percent overlap of streamlines, from different ROIs, into the same striatum voxel.

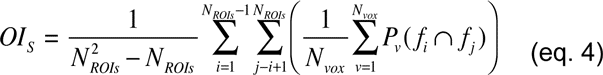

For all *N_vox_* voxels in the striatum mask, the conditional probability, *P_v_* (*f_i_* ∩ *f_j_*), that at least one streamline from any pair of cortical ROIs, *f_i_* and *f_i_*, terminates within the voxel was determined. If an overlap of streamlines was detected, then *P_v_* (*f_i_* ∩ *f_j_*) = 1, otherwise *P_v_* (*f_i_* ∩ *f_j_*) = 0. This was averaged across all striatal voxels to provide a probability of overlap of projections for any given pair of cortical ROIs. This process was repeated for all pairs of cortical ROIs and averaged to create a composite index for each subject. The OI was calculated independently for the left and right hemispheres networks.

## Results

### Networks associated with response conflict

Behavioral responses in the scanner were consistent with standard conflict resolution and adaptation effects in the Stroop task (Stroop, 1935; Macleod, 1991; Gratton et al., 1992; Botvinick et al., 2001). On all correct trials, there was a main effect of stimulus condition on response times (RT; Fig. 1A; F(58,2) = 67.88, p < 0.001). Compared to neutral control trials, subjects were slower to respond when there was a cue conflict, (incongruent trials; t(27) = 10.02, p < 0.0001), and faster on trials with redundant cues, (congruent trials; t(27) = −5.68, p < 0.0001). Responses during incongruent trials were modulated depending on the structure of the previous trial. When the previous trial was also incongruent participants were 32 ms faster to respond than when the previous trial was congruent (Fig. 1B; t(27) = 2.82, p = 0.004). Participants were slightly more accurate (2% fewer errors) on incongruent trials when the previous trial was also incongruent (Fig 1C; t(27) = 2.05, p = 0.025). A similar repetition effect was observed on congruent trials, but attenuated. On average, participants were 19 ms faster on a congruent trial when the previous trial was also congruent, rather than when the previous trial was incongruent (Fig 1B; t(27) = −11.22, p < 0.0001). No advantage was observed for repeating congruent trials on the error rate (t(27) = 0.41, p = 0.34), however, this may be due to a low base rate of errors on congruent trials to begin with (∼1%).

**Figure 1.**
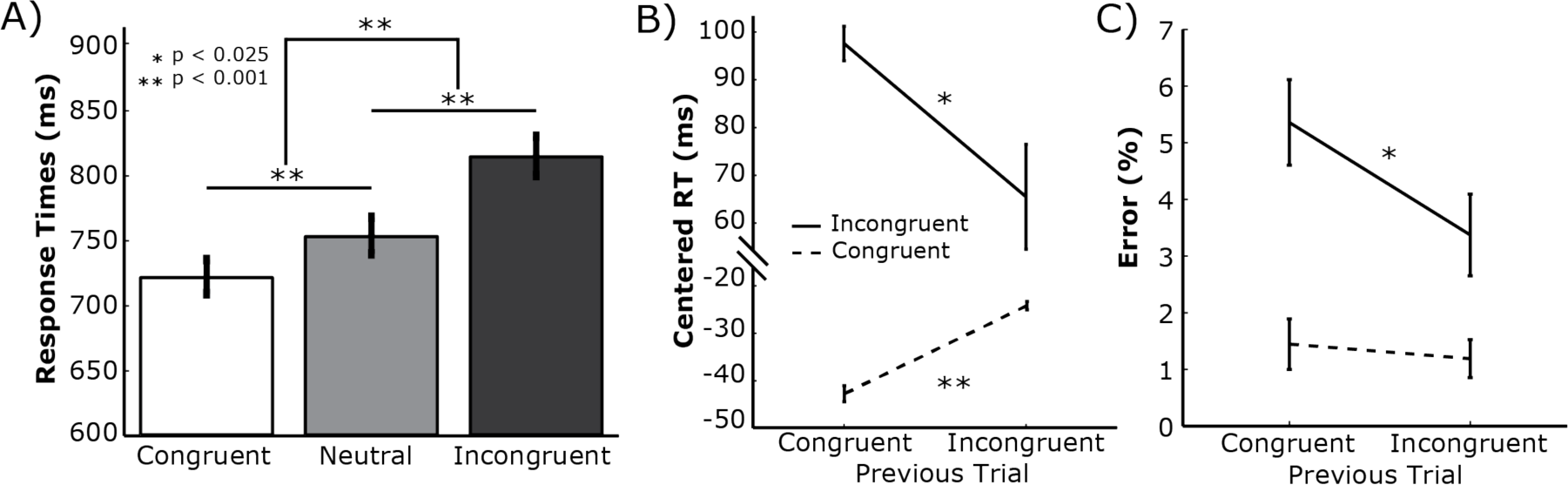
Behavioral responses during the color-word Stroop task. A) In general, response times (RT) on accurate Incongruent trials were slower than Neutral trials while RTs for Congruent trials were faster than Neutral trials. B) Incongruent trials (solid black line) preceded by Incongruent trials had faster responses than when they were preceded by a Congruent stimulus. In contrast, Congruent trials that were preceded by an Incongruent stimulus were slower when two Congruent trials were repeated. All RTs were mean centered prior to analysis. C) The advantage for condition repetition was also seen in accuracy, with repetition of Incongruent trials leading to fewer errors on the second trial. This effect, however, was not seen for Congruent trials. All error bars show standard error across subjects.

In order for conflict adaptation to occur, a critical computation has to occur on or immediately after the preceding trial. To isolate regions associated with cue incongruency, a whole-brain random effects analysis was performed to identify regions selectively responsive to incongruent trials, compared to neutral trials. Twenty distributed regions were associated with task-specific responses during Incongruent trials, when compared against neutral trials (Fig. 2A–B, Table 1). Many of these have been previously associated with incongruency in the Stroop task (Banich et al., 2000), as well as other cue conflicts in other tasks (Casey et al., 2000; Weissman and Carp, 2013).

**Figure 2.**
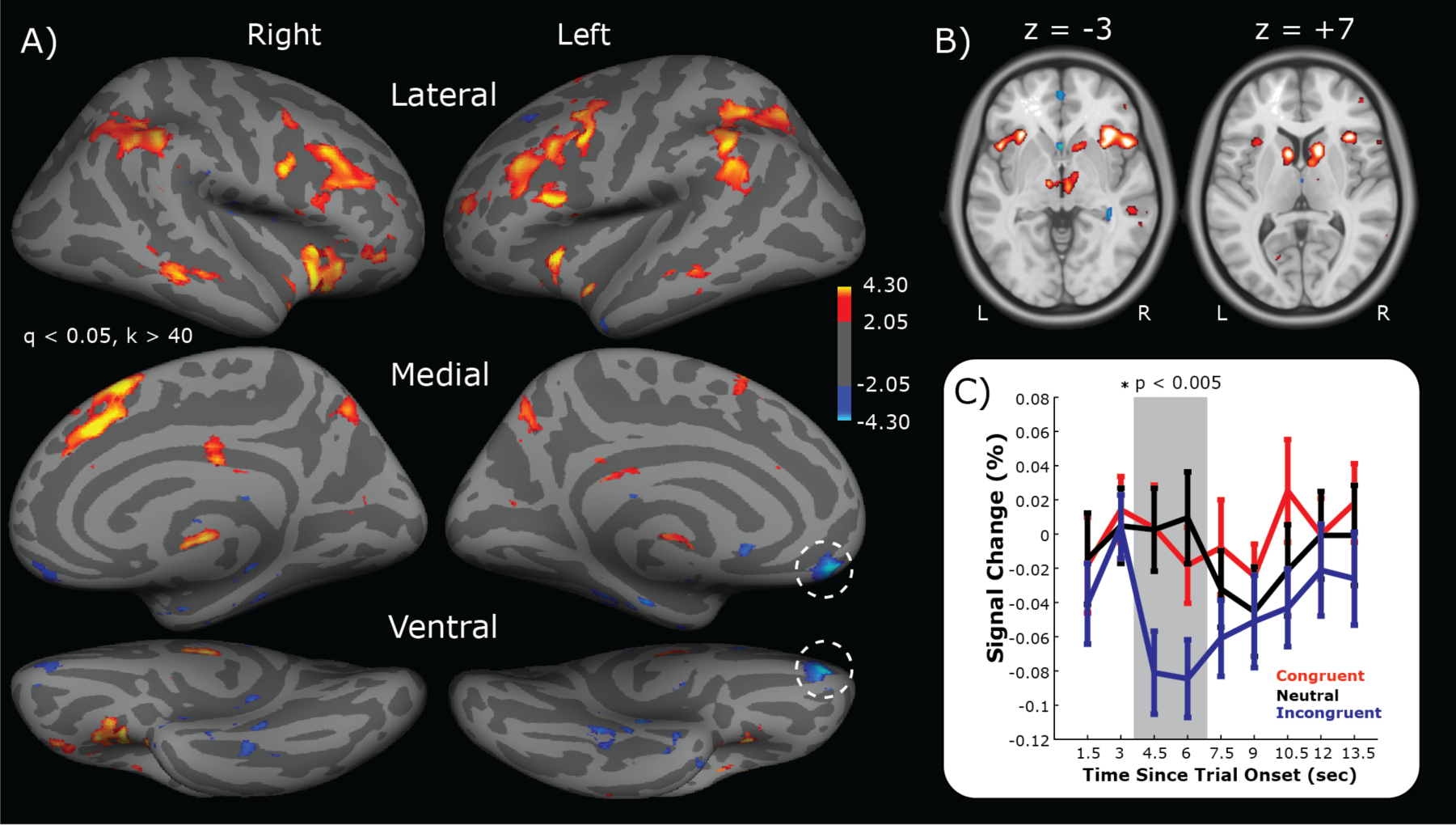
Stroop related BOLD responses. A) Activation maps on an inflated brain showing voxels with differential BOLD responses to cue incongruency (i.e., Incongruent vs. Neutral trials). Warm colors indicate areas where the Incongruent trial response was more positive than the neutral trial response. Cool colors indicate areas where the neutral trial response was more positive. Activation maps are adjusted for multiple comparisons using a false discovery rate (q) of less than 0.05 and restricted to clusters of 40 or more continuously connected voxels. Dashed circles highlight the negative cluster on the gyrus rectus of the medial orbitofrontal cortex (mOFC). B) Two axial slices showing the same activation patterns as in A. Slice position in MNI coordinates are shown above each slice. C) Trial-locked BOLD responses from the mOFC cluster for each task condition. Averaged signal change values in the peak of the BOLD response (gray bar) show a significant modulation based on condition type (repeated measures F(2,54) = 6.61, p = 0.003). Error bars show standard error across subjects.

**Table 1.** Clusters with Incongruent-related responses, compared to Neutral trials. MNI coordinates, t-statistic and p-values are for the peak voxel in the cluster. intrapareital sulcus, IPS; middle frontal gyrus, MFG; temporal-parietal junction, TPJ; subthalamic nucleus, STN; precentral sulcus, PCS; supramarginal gyrus, SMG; middle temporal gyrus, MTG; parahippocampal cortex, PHC.

| ROI | NVox | MNI Coordinates (x,y,z) | Peak T | Peak P |
| --- | --- | --- | --- | --- |
| L_Caudate | 93 | -10, 10, 8 | 5.55 | p<0.0001 |
| L_IPS | 712 | -54, -44, 38 | 5.69 | p<0.0001 |
| L_Insula | 267 | -32, 22, 0 | 7.11 | p<0.0001 |
| L_MFG | 842 | -48, 18, 28 | 7.77 | p<0.0001 |
| L_MTG | 41 | -56, -34, -8 | 4.86 | p<0.0001 |
| L_PHC | 56 | -14, -12, -18 | -3.62 | 0.0006 |
| L_mOFC | 87 | -10, 52, -16 | -3.61 | 0.0006 |
| L_TPJ | 64 | -58, -50, 28 | 4.70 | p<0.0001 |
| L_STN | 26 | -10, -14, -2 | 5.06 | p<0.0001 |
| Precuneus | 98 | 8, -68, 44 | 4.35 | p<0.0001 |
| R_Medial SFG | 819 | 6, 16, 60 | 6.70 | p<0.0001 |
| R_Caudate | 223 | 14, 12, 4 | 6.06 | p<0.0001 |
| R_MFG (ventral) | 45 | 48, 40, -8 | 4.94 | p<0.0001 |
| R_IPS | 716 | 58, -54, 40 | 5.04 | p<0.0001 |
| R_Insula | 742 | 30, 18, -12 | 8.26 | p<0.0001 |
| R_MFG | 736 | 50, 16, 28 | 6.21 | p<0.0001 |
| R_MTG | 68 | 64, -30, -10 | 4.86 | p<0.0001 |
| R_PCS | 97 | 48, 12, 54 | 5.69 | p<0.0001 |
| R_STN | 73 | 6, -16, -2 | 5.15 | p<0.0001 |
| R_SMG | 52 | 62, -48, 20 | 4.93 | p<0.0001 |

Previous reports of reward contingency and reward sensitivity biasing the degree of conflict adaptation (van Steenbergen et al., 2009; Braem et al., 2012) would suggest that incongruency may also be reflected in the responses of areas also linked to reward processing. Consistent with this idea a cluster on the gyrus rectus of the medial orbitofrontal cortex (mOFC) was identified that exhibited selective responses for incongruent trials, relative to neutral trials (dashed circles in Figure 2A). This mOFC cluster was one of two clusters identified in the whole-brain analysis that had a negative contrast value, i.e., more positive BOLD responses during incongruent trials than neutral trials (see Table 1). To better understand the nature of this negative contrast the average trial-evoked BOLD response for each stimulus condition was estimated (Fig. 2C). Unlike congruent and neutral trials, the mOFC exhibited a clear time-locked response to incongruent trials, reflected as a dip in the BOLD signal. Therefore, this region was responsive to conflict in response cues, but with a negatively directed evoked BOLD response.

In order to understand how activity at the various regions of this distributed network of incongruency-related areas might relate to trial-by-trial behavioral performance, a single-trial regression analysis was adopted to measure of evoked BOLD responses on individual trials (see Methods). This trial-by-trial BOLD response was then correlated with RTs, across all conditions, for each incongruency-related cluster (Fig. 3A). Of the 20 ROIs associated with incongruent trials, trial-wise BOLD responses in 15 regions were significantly correlated with RTs on the current trial. However, the mOFC cluster was not associated with current trial RTs. But as hypothesized, the evoked BOLD signals in the gyrus rectus were correlated with RT on the following trial (Fig 3B), suggesting that the mOFC may be playing a role in updating future responses based on current trial outcomes.

**Figure 3.**
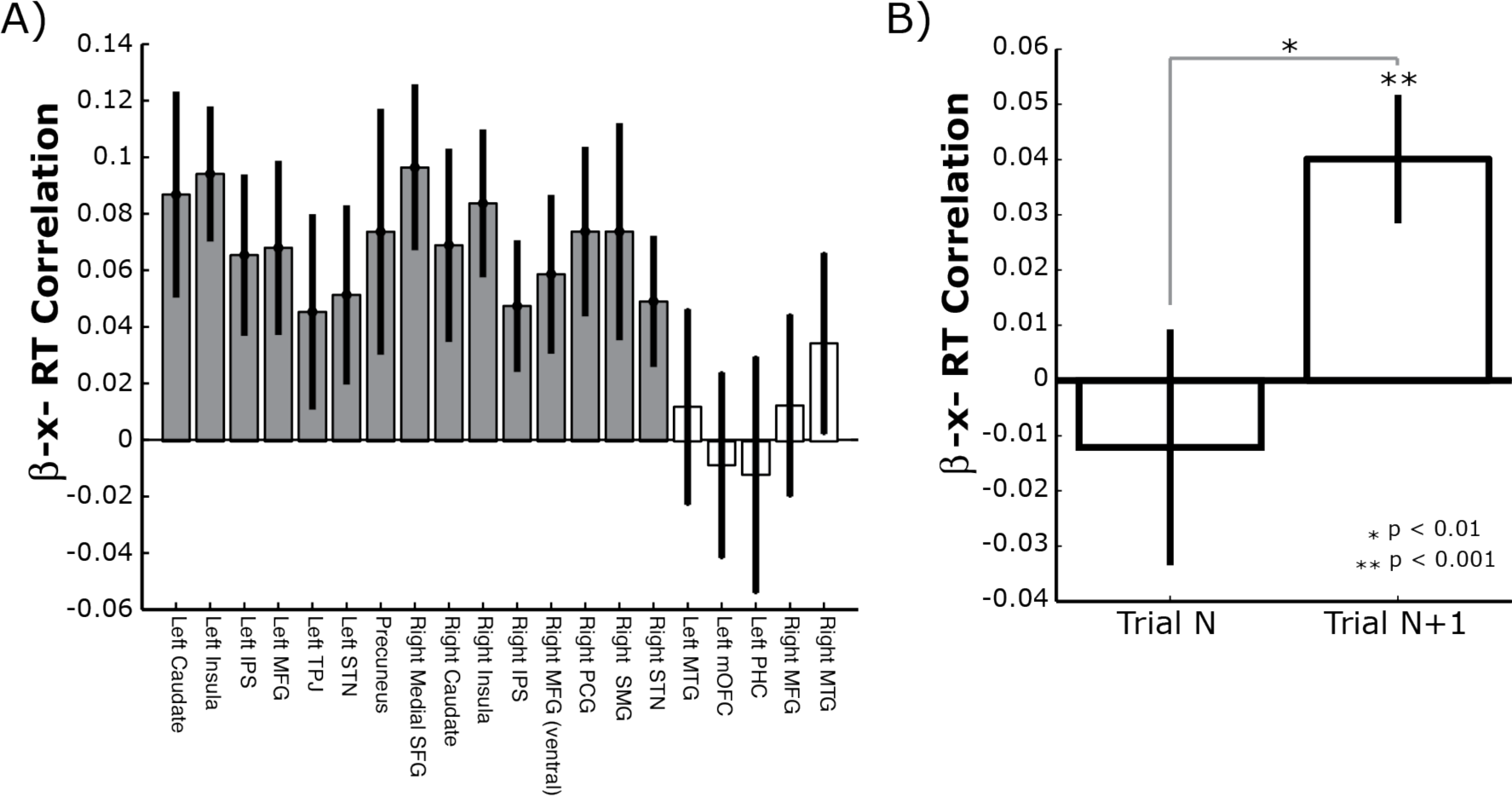
Brain-behavior correlations. A) Correlation coefficients for the association between RT on the current trial and the single-trial BOLD estimate from 20 regions of interest (ROIs). Dark bars indicate ROIs with significant trial-wise associations across subjects (FDR corrected, q < 0.05). Error bars show the Bonferroni corrected 95% confidence interval (CI) across subjects. Same abbreviation convention as Table 1. B) Correlation for the mOFC responses against current trial (Trial N) RT, showing the same data as in A, and the following trial RT (Trial N + 1). Same plotting convention as A.

### Distributed networks linking orbitofrontal responses to behavior

At first glance, seeing that mOFC activity was not correlated with current trial performance may appear contradictory to the hypothesis that ventral corticostriatal systems are involved in the process of conflict adaptation. After all, according to neural theories of reinforcement learning, an evaluation of current trial outcomes is necessary to modify future responses (Dayan and Abbott, 2001). However, given the large and distributed network of areas associated with incongruency in the previous analysis it is possible that a residual association between mOFC and current trial RT is masked by indirect pathways linking orbitofrontal responses to current trial outcomes.

To test this hypothesis, a conditional regression analysis was used to identify indirect pathways linking gyrus rectus activity to current trial responses (see Methods; (Preacher and Hayes, 2008a)). This analysis revealed three bilateral clusters that were indirect pathways linking mOFC responses to RT on the current trial (Fig 4A): the middle frontal gyrus, the insula, and caudate nucleus. Thus there were dorsolateral, ventrolateral and striatal regions that statistically mediated the association between mOFC responses on the current trial and the speed of behavioral responses on that same trial.

**Figure 4.**
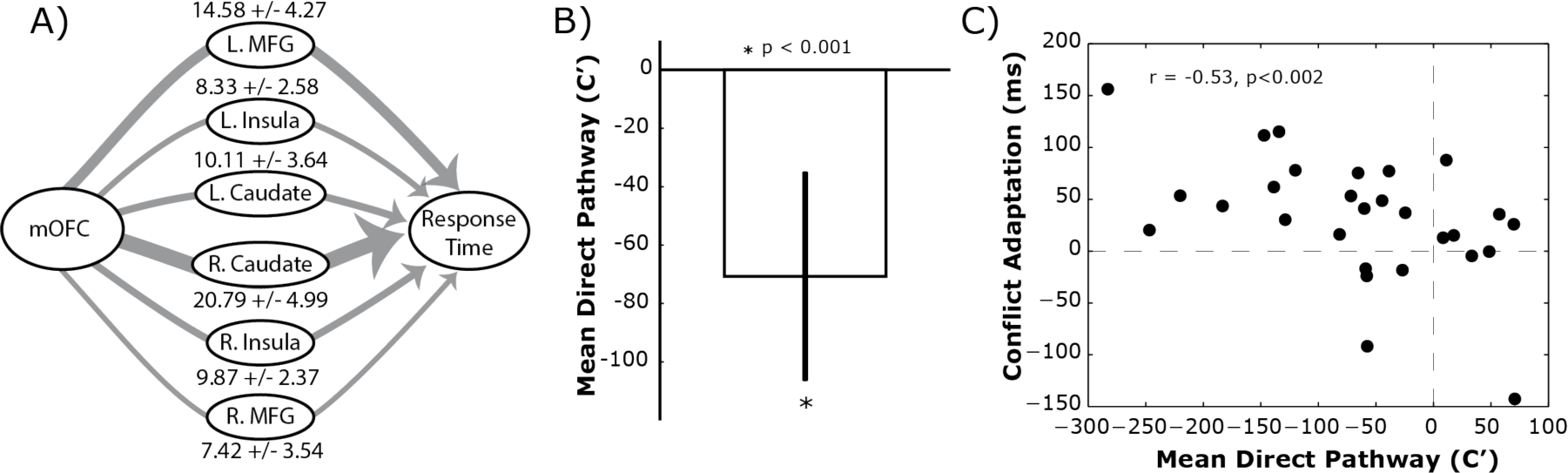
Mediating network pathways. A) Statistical mediation analysis revealed three bilateral regions that served as indirect pathways mediating the association between mOFC responses and RT on the same trial. Thickness of the lines illustrates the magnitude of the indirect pathway coefficients (a*b). Mean indirect pathway coefficient and standard deviation, across subjects, are shown above each mediating ROI. B) Recovered direct pathway (C’) coefficient, showing the residual relationship between mOFC and current trial RT after accounting for indirect pathways. Same plotting conventions as used in Figure 2. C) Individual differences analysis showing the association between a subject’s direct pathway magnitude and the degree of conflict adaptation.

By nature of the conditional regression analysis (see eq. 3 in Materials and Methods), the direct pathway (c’) represents the residual relationship between mOFC responses and RT, after accounting for the indirect interactions with the mediating ROIs: i.e., *c*′ *X* = *Y* − *bM* − *η*. Averaged across all significant indirect pathways, there was a consistent negative c’ pathway between mOFC and behavior (Fig. 4B; t(27) = −3.90, p = 0.0002); Fig 3B). This effect was also significant for the residual from each indirect pathway when looked at individually (all p’s < 0.0002). Taking into account the direction of the evoked BOLD response in the gyrus rectus (Fig. 2C), this means that stronger (i.e., more negative) evoked responses in the mOFC are associated with slower reaction times on the current trial, after controlling for indirect associations with the lateral prefrontal cortex, caudate nucleus and insula. Thus the mOFC-RT association on the current trial is normally obscured by the interactions between the indirect regions that also relate to both mOFC responses and behavior.

In order to understand what, if any, predictive value this direct pathway has on conflict adaptation, an individual differences analysis was performed between the direct pathway coefficients (Fig. 4B) and conflict adaptation scores (see *Behavioral Analysis Section* and Fig 1B). The average direct pathway coefficient (*c’*), across all significant indirect paths, was negatively correlated with the degree of conflict adaptation (Fig. 4C; r = −0.53, p = 0.001). Because there is a separate *c’* coefficient estimated for each indirect pathway, it is possible to isolate which mediating paths are accounting for the most variance in the mOFC-RT relationship. After controlling for multiple comparisons (q < 0.05), the residual *c’* coefficients from the bilateral lateral prefrontal (left: r = −0.55, p < 0.001; right: r = −0.53, p < 0.001) and bilateral caudate (left: r = −0.51, p < 0.001; right: r = −0.54, p = 0.001) pathways were also significantly associated with conflict adaptation magnitude. The direction of these relationships implies that the subjects with stronger (i.e., more negative) *c’* coefficients had the greatest adaptation effect on subsequent trials. More importantly, however, none of the indirect (*a*b*) pathways themselves were associated with the magnitude of conflict adaptation (all p’s > 0.15).

These functional imaging results reveal that the mOFC activity correlates with the speed of future responses and is indirectly associated to current trial RT via mediating pathways in lateral frontal and striatal regions. Most importantly, however, after accounting for these indirect pathways, the strength of the residual mOFC-RT relationship on the current trial predicts the degree of conflict adaptation. Taken together, these findings are all consistent with the hypothesis that the ventral corticostriatal network is associated with trial-by-trial adaptation in cue-conflict tasks.

### Convergence of striatal inputs

One common connectivity pattern links all the regions shown in Figure 4A: the orbitofrontal and lateral prefrontal regions all send feedforward projections into the caudate nucleus (Haber and Knutson, 2010). Thus, anatomically speaking, the striatum may represent a central integration point during the response selection and adaptation processes.

To evaluate this anatomical hypothesis, fiber tractography data from diffusion spectrum imaging was used to map out the underlying white matter pathways from twelve frontal regions and one subcortical source (see Methods). Figure 5A,B shows an example tractography run from a single subject. Streamlines from orbitofrontal (red), medial wall (green), lateral frontal (yellow), insula (cyan) and thalamus (violet) all terminate cleanly within the mask for the caudate nucleus and putamen (black region of interest). Figure 5B shows the consistent topography of streamline endpoints, with orbitofrontal fibers primarily terminating ventrally from medial and lateral prefrontal streamlines.

**Figure 5.**
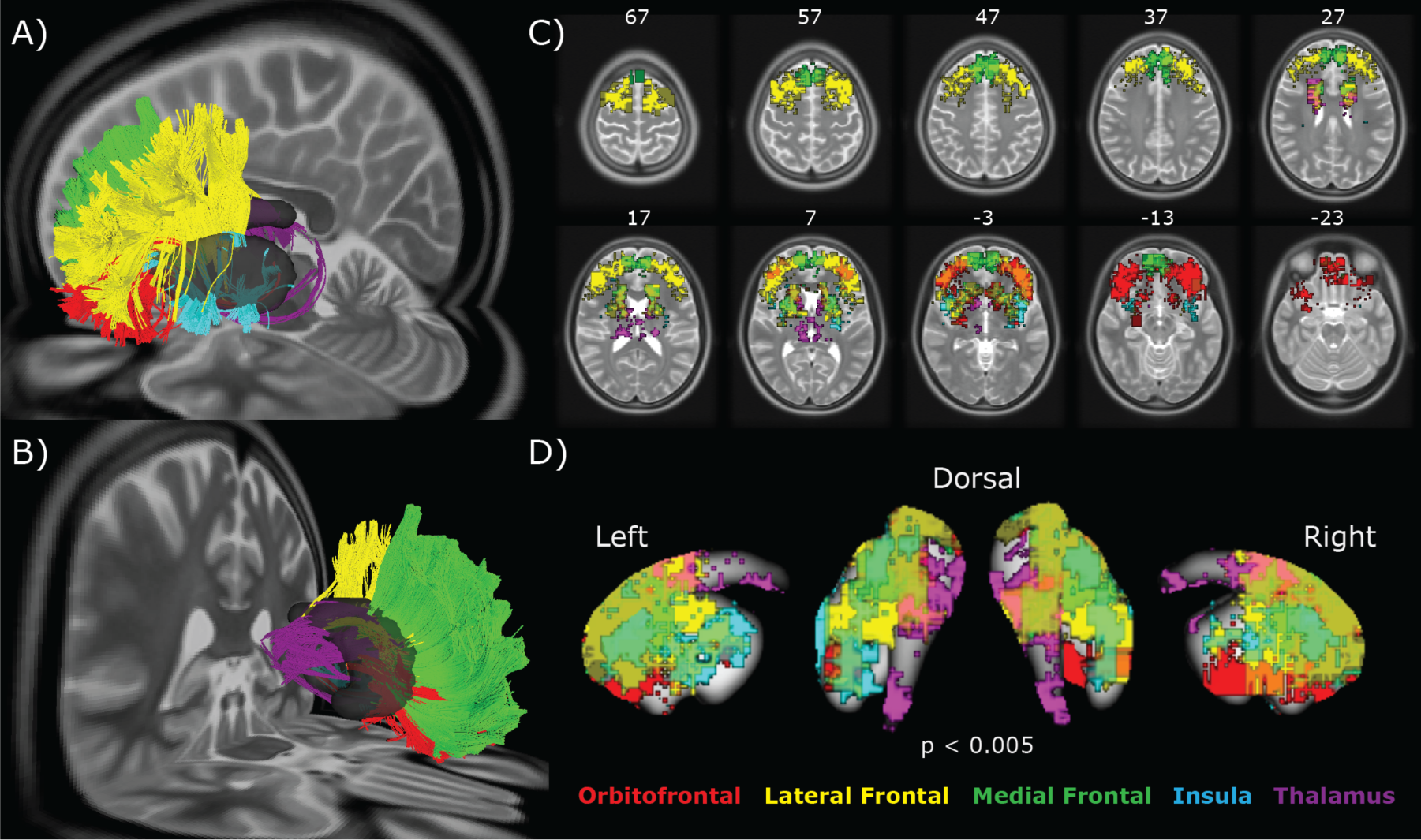
Topography of corticostriatal projections. A,B) Example deterministic tractography results from a single subject showing tracked left hemisphere projections only. Streamlines are colored based on cortical grouping. *A* shows a lateral view while *B* shows a medial view. C) Voxel-wise maps showing the locations of highest endpoint density of corticostriatal projections, across subjects. Voxels are thresholded at a t > 2.75 and p < 0.005, uncorrected. D) Same data as shown in C, projected on a template of the striatal nuclei.

In order to quantify this topography at the group level, the endpoint location density of streamlines was averaged across subject for each ROI and voxels with consistent streamline terminations were determined using a one-sampled t-test. Figure 5C shows the cortical and subcortical terminations fields, across subjects, overlaid on a T2 anatomical template.

Generally there was a gross segmentation of endpoint fields within the striatum that is consistent with previously reported topographies of inputs from frontal areas (Draganski et al., 2008; Haber and Knutson, 2010; Verstynen et al., 2012). For example, orbitofrontal projections tended to terminate in the rostral and ventral aspects of the striatum, while lateral prefrontal regions terminated more dorsal and caudal regions in the body of the caudate. Yet there is also substantial overlap in these endpoint fields along the striatum. This is more clearly seen in the close up of the striatum in Figure 5D. In general, the streamlines from medial, lateral and orbital sources shared a moderate degree of overlap in the rostral striatum, particularly the near the shell of the striatum. In fact, the cluster of caudate activity observed in the fMRI results (Fig 2B) appears to be situated just between projection fields from all areas tracked, except for the insula, in the rostral caudate. Although given the difference in distortion between functional EPI and DWI sequences, it is difficult to attribute the caudate activation to specific inputs from any cortical area.

This overlap of streamline endpoints into the rostral aspects of the caudate is consistent with previous tracer studies in non-human primates (Haber et al., 2006). Specifically, Haber and colleagues (2006) propose that integration in the striatum happens when diffuse (i.e., low density) projections from one cortical area overlap with focal (i.e., high density) projection fields from another cortical area. Anatomically, the current results show evidence of these diffuse and focal projection field overlaps. Figure 6A shows the mapped white matter projections from the middle frontal gyrus (purple streamlines) and medial orbitofrontal gyrus (cyan streamlines) for a single subject. The start and end locations of these streamlines are shown in Figure 6B, with the caudate and putamen regions of interest shown in black. As is highlighted in a close-up inset image (Figure 6C), there are several regions of overlap between the two streamline sets (dashed circles).

**Figure 6.**
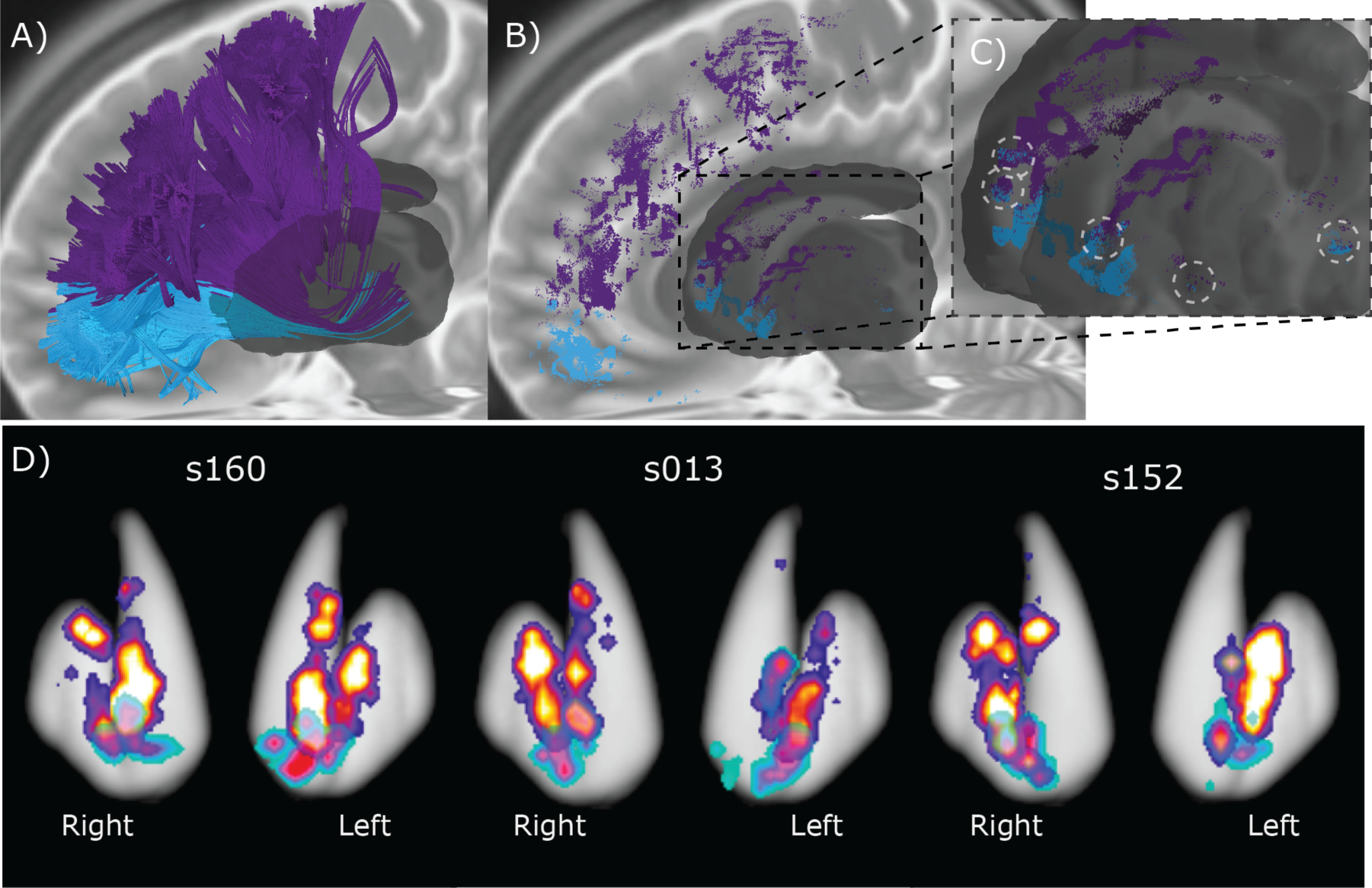
Overlapping corticostriatal projections. A) Streamlines from two ROIs (middle frontal gyrus, purple; medial orbitofrontal gyrus, cyan) from a different subject than shown in Fig. 5A,B. B) Streamline endpoint locations for same data shown in A. Striatum ROI mask shown in dark gray. C) Close up of endpoint locations along the striatum. Dashed circles highlight regions where adjacent streamlines overlap. D) Density maps of streamline endpoints, for the same two pathways shown in A, at each voxel along the striatum for three subjects. Subject s160 is the subject shown in A. Warmer maps show endpoint densities from middle frontal gyrus. Cooler maps show endpoint densities from medial orbitofrontal gyrus.

To get a better handle on the degree of overlap and the density of the projection fields, the endpoint densities along the striatum were calculated for each voxel in the same two pathways. Figure 6D shows these densities in three example subjects. Subject s160 is the same subject as is shown in Figure 6A–C (i.e., showing the same data sets). Consistent with the diffuse overlap hypothesis (Haber and Knutson, 2010), the centers of mass for the two projection fields do not overlap; however, there is consistent overlap of the less dense portions of the projection fields in all three subjects.

The degree of overlapping fiber streamlines from adjacent cortical regions to the same striatal voxels (see Methods) was next quantified using an Overlap Index score (OI; see Methods). The OI defines the percent of the time that any pair of region of interest overlap on voxels in the mask of the striatal nuclei. This score was averaged across all voxels in the left and right striatum masks separately, providing a composite index for the level of local overlap for each subject and hemisphere. Overall, there was a greater degree of overlap in the right hemisphere (0.40, 95% confidence interval upper bound = 0.50, lower bound = 0.29) than the left hemisphere (0.22, 95% CI upper bound = 0.29, lower bound = 0.15). It should be noted, however, that even though striatal spiny neurons are known to receive a high concentration of convergent inputs (Kincaid et al., 1998), this score cannot be interpreted as capturing shared collaterals on the same striatal neurons as this is beyond the spatial resolution of current diffusion imaging methods. Instead, this OI reflects the proximity of inputs from different anatomically defined cortical regions.

The functional significance of this structural measure was assessed using a correlation analysis between the OI and the indirect (a*b) and direct (c’) pathways from the mediation model. While structural overlap did not correlate with the indirect pathways (all p’s > 0.39), the overlap of corticostriatal projections in the left hemisphere was negatively correlated with the magnitude of the direct pathway coefficient (Fig. 7A; r = −0.36, p = 0.032). The direction of this relationship implies that subjects with a greater degree of overlap of anatomical connections into the striatum also had stronger conditional mOFC-RT relationships. This pattern was not observed in the right hemisphere (Fig. 7B; r = 0.13, p = 0.22); however, this laterality may be due to the fact that only the left mOFC was included in the calculation of the direct pathway coefficients. Nonetheless, the pathway with the most predictive value for conflict adaptation was itself correlated with local overlap of frontal corticostriatal projections, highlighting a possible structural mechanism for the integration of executive and reward processes during response updating.

**Figure 7.**
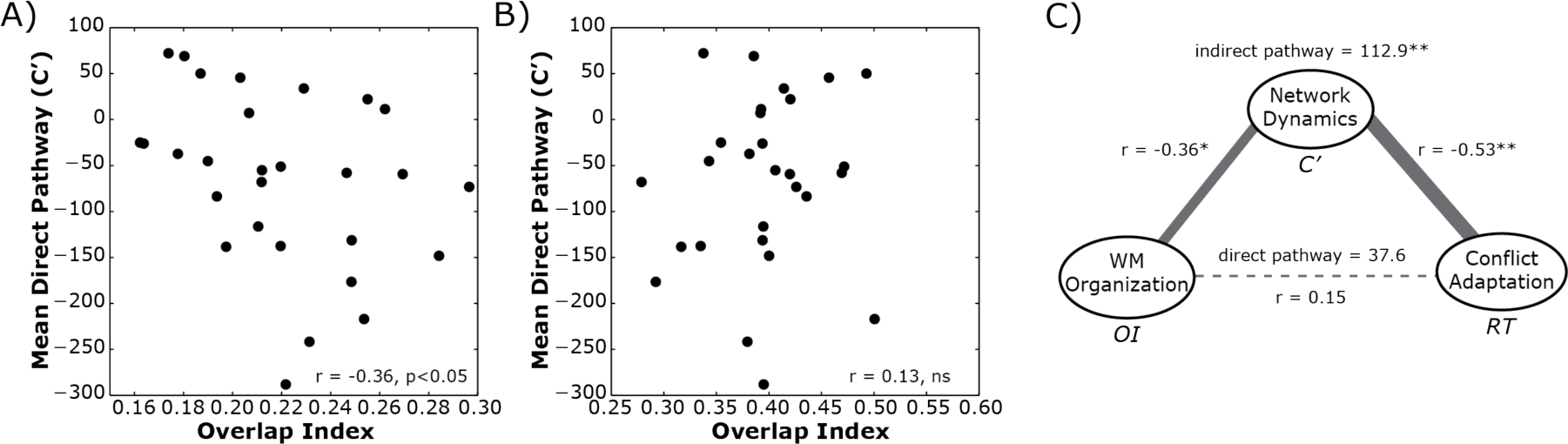
Structure-function associations. A,B) Correlations between direct pathway coefficients from the functional analysis (see Fig. 4) and degree of structural overlap in the fiber tractography analysis for the left hemisphere, A, and right hemisphere, B, pathways. C) Full structure-function-behavior model showing the associations between structural overlap in the white matter (WM) pathways (OI Score), functional brain-behavior links during the Stroop task (C’ score from model in Figure 4A–B) and behavioral conflict adaptation scores (RT). Data for left hemisphere pathways only. Asterisks indicate significant associations at p < 0.05 (*) and p < 0.002 (**).

### A structure-function-behavior pathway

The preceding analysis revealed that subjects with a greater degree of overlapping anatomical connections into the striatum also have stronger, i.e., more negative, conditioned associations between mOFC responses and current trial RT. Furthermore, stronger conditional mOFC-RT relationships correlated more with the degree of conflict adaptation on future incongruent trials. Taken together, these associations hint at the possibility that individual differences in the structural topography of corticostriatal networks may provide a computational constraint on the process of behavioral adaptation. In support of this hypothesis, the conditional mOFC-RT effect was found to be a significant indirect pathway linking structural overlap scores and conflict adaptation (a*b = 112.4, 95% CI = 44.0–183.0, p < 0.001; Figure 7C). This indirect pathway association was significant despite the fact that the simple correlation between OI and conflict adaptation correlation was not significant (r = 0.15, p = 0.23). Accounting for the indirect associations via the functional network dynamics, also did not yield a significant the direct path between OI and conflict adaptation (c ’ = 37.6, 95% CI = −42.4–131.4, p = 0.19), suggesting any relationships white matter had to behavior were fully dependent on indirect associations with functional processes. Taken together, these patterns of associations suggest that structural overlap of corticostriatal projections has a conditional relationship with behavioral adaptation through functional network dynamics that bind mOFC activity to response times.

## Discussion

Using a standard cue conflict paradigm, the present study revealed that responses in a medial region of the orbitofrontal cortex, on the gyrus rectus, are strongly associated with the speed of upcoming decisions through interactions with frontal and striatal regions. In particular, stronger (i.e., more negative) evoked responses in the mOFC correlated with faster RTs on the following trial and, after accounting for interactions with lateral prefrontal areas and the caudate nucleus, slower behavioral responses on the current trial. The stronger the relationship between mOFC and current trial RT, the better a subject adapted to conflicting response cues. The fact that these brain-behavior associations with mOFC were conditioned on concurrent responses in lateral prefrontal cortex and caudate nucleus suggests that any contribution the mOFC has to response adaptation depends on an integration of information from multiple cortical and subcortical sources. One possible mechanism for this integration is via convergent corticostriatal inputs (Haber et al., 2006). Indeed, analysis of structural connections within rostral corticostriatal pathways confirmed that variation in the amount of overlapping projections from adjacent cortical sources into the striatum predicted the efficiency of the functional brain-behavior relationships: i.e., more overlap of white matter projections into the striatum correlated with stronger mOFC-RT associations that, in turn, correlated with greater conflict adaptation.

The particular cluster of task-related voxels in the mOFC was found on a rostral aspect of the gyrus rectus, which is sometimes referred to as the straight gyrus. The gyrus rectus is part of Brodmann’s area 11. It is medial to the olfactory sulcus and lateral to the longitudinal fissure. Although very little is known about its functional role in behavior, it is sometimes considered part of the orbitofrontal complex because of its response properties and clinical symptomatology. In a meta-analysis of 142 imaging studies, Liu and colleagues (2010) reported that the most medial and rostral aspects of the orbitofrontal cortex, including portions of the gyrus rectus, were most strongly associated with positive reward experiences and variation in reward outcome, as opposed to anticipation or evaluation (Liu et al., 2011). This would be consistent with the present study and with neural network models of the orbitofrontal cortex in reinforcement learning (Frank and Claus, 2006; Ratcliff and Frank, 2012). Clinically, lesions to the medial orbifrontal cortex, including the gyrus rectus, are most often associated with disinhibition syndromes (Cummings, 1998; Hornak et al., 2003); however, these lesions often include significant potions of the medial orbitofrontal gyrus or sections of the anterior cingulate cortex, making it difficult to infer behavioral patterns that are specific to gyrus rectus damage itself. In fact, the single reported case study of a focal lesion restricted to the left gyrus rectus found that the patient did not present with disinhibition symptoms, but instead exhibited athymhormic syndrome, which is a rare syndrome characterized by blunted affect and extreme apathy (Winckler et al., 2011). Given the limited understanding as to the function of the gyrus rectus, the current findings linking it to rapid trial-by-trial updating may help to shed some light on the particular cognitive roles this region performs.

Perhaps the most striking observation of the present study is the association between mOFC activity and future behavioral responses in a task without explicit reward contingencies. Indeed, it is well known that the orbitofrontal cortex is critical for flexible behavior in reinforcement learning contexts, particularly in the face of quickly changing reward contingencies (see (Schoenbaum et al., 2010)). Mechanistically it has been proposed that the orbitofrontal cortex helps to modify future behavior by rapidly learning new associations between cues and outcomes through indirect associations with other brain areas (Rolls et al., 1996; Frank and Claus, 2006). In this way the orbitofrontal cortex is thought to learn to predict outcome expectancies, including but not limited to predicting expected rewards (Schoenbaum et al., 2010). This idea is also consistent with ‘actor-critic’ models of ventral striatal areas during reinforcement learning, in which the striatum contributes to the estimation of internal state-values that are compared against incoming sensory information on future trials in order to generate appropriate error signals (O’Doherty et al., 2004).

Together, these two state-updating models of ventral corticostriatal networks are consistent with the pattern of results found in the current study. Greater monitoring of stimulus features that lead to correct responses (i.e., stronger mOFC response) during conflict trials (i.e., when RTs are slower) increases the gain of attending to those features in the immediate future (i.e., faster RTs on the next trial). In this way the orbitofrontal cortex acts as a modulator of executive decisions relayed through the prefrontal corticostriatal circuits that make the final response choice, rather than a mediator of conflict adaptation itself. This idea is consistent with neural network models of fast reinforcement learning (Frank and Claus, 2006; Ratcliff and Frank, 2012) where orbitofrontal projections modulate the sensitivity of go/no-go pathways within the basal ganglia that are triggered by prefrontal selection processes. The observation in the current study that lateral prefrontal and striatal regions served as indirect pathways linking orbitofrontal activity to behavioral responses is consistent with this integration and modulation model.

Mechanistically, this orbitofrontal priming of the indirect (go) and direct (no-go) pathways in the basal ganglia requires an integration of information between ventral and dorsal corticostriatal circuits. Although traditionally viewed as completely parallel and independent systems (Alexander et al., 1986), it is becoming increasingly apparent that there is a moderate degree of integration across neighboring basal ganglia loops. In frontal cortico-basal ganglia systems, Haber and Knutson (2010) proposed three types of integration pathways that would allow for convergence of executive control and reward information within the basal ganglia: feedback loops relaying information from ventral striatum to the dorsal striatum via substantia nigra dopamine pathways, convergence of feedback thalamicocortical projections, and local overlap of corticostriatal inputs themselves. This last integration mechanism stems largely from neuroanatomical observations in animals, where diffuse projections from one cortical area terminate in regions of the striatum that contain more dense projections from another cortical origin (Haber et al., 2006). In humans a similar pattern of diffuse projections was observed in corticostriatal pathways that were tracked using fiber tractography on diffusion weighted imaging data (Verstynen et al., 2012). This study found an asymmetry in the direction of these diffuse projections, with more streamlines starting in rostral frontal areas and terminating in more caudal striatal regions than vice versa. This means that there were more projections from orbitofrontal areas that terminated in the dorsal and caudal striatum than there were projections from dorsolateral prefrontal cortex that terminated in the rostral and ventral striatum. When considered within the context of the present study, the direction of this asymmetry suggests that this overlap of corticostriatal inputs may be a mechanism of information convergence that is relevant for conflict adaptation. Indeed, the current study found that individual differences in overlapping corticostriatal projections predicted individual differences in both functional dynamics and brain-behavior relationships during conflict adaptation. This strongly suggests that integration of modulatory signals from orbitofrontal cortex and executive control signals from lateral prefrontal cortex happens, at least in part, through common inputs in the striatum. It is entirely possible that reciprocal loops with midbrain areas and convergent feedback projections to the thalamus also predict conflict adaptation dynamics; however, visualizing these connections is beyond the capabilities of current white matter visualization methods in humans.

It is perhaps curious that the present study did not find an association between activity in medial wall areas, like the anterior cingulate cortex, and conflict adaptation (Sheth et al., 2011, 2012; Kim et al., 2013a, 2013b). As was mentioned at the beginning of this article, the behavioral phenomenon of conflict adaptation actually reflects an amalgamation of many different cognitive processes, including contingency learning, feature integration, congruency sequencing effects (Schmidt, 2013a, 2013b) and reinforcement learning (van Steenbergen et al., 2009; Braem et al., 2012). It is possible that ventral striatal updating processes have a larger effect size than conflict monitoring processes, meaning that the current experimental design is underpowered to reveal any brain-behavior relationships between medial wall areas and conflict adaptation on a trial-by-trial basis. Within the standard Stroop task, it is not possible to isolate the pure congruency sequencing effects thought to be mediated by conflict monitoring regions like the anterior cingulate cortex (Schmidt, 2013a, 2013b). Explicitly dissociating the different cognitive components that underlie conflict adaptation, along with their mediating neural systems, will be important directions for future research.

## Acknowledgements

This research was sponsored in part from PA Department of Health Formula Award #SAP4100062201 and by the Army Research Laboratory under Cooperative Agreement Number W911NF-10-2-0022. The views and conclusions contained in this document are those of the authors and should not be interpreted as representing the official policies, either expressed or implies, of the Army Research Laboratory or the U.S. Government. The U.S. Government is authorized to reproduce and distribute reprints for the Government purposes not withstanding any copyright notation herein. The author would like to thank Daniel Weissman and Jean Vettel for their helpful comments on the initial phases of this study and Kevin Jarbo, Patrick Buekema, and Fang-Cheng Yeh for their critiques on early versions of the manuscript.

